## Supplementary material for "Cholangiocytes contribute to hepatocyte regeneration after partial liver injury during growth spurt in zebrafish": Figures S1-S8

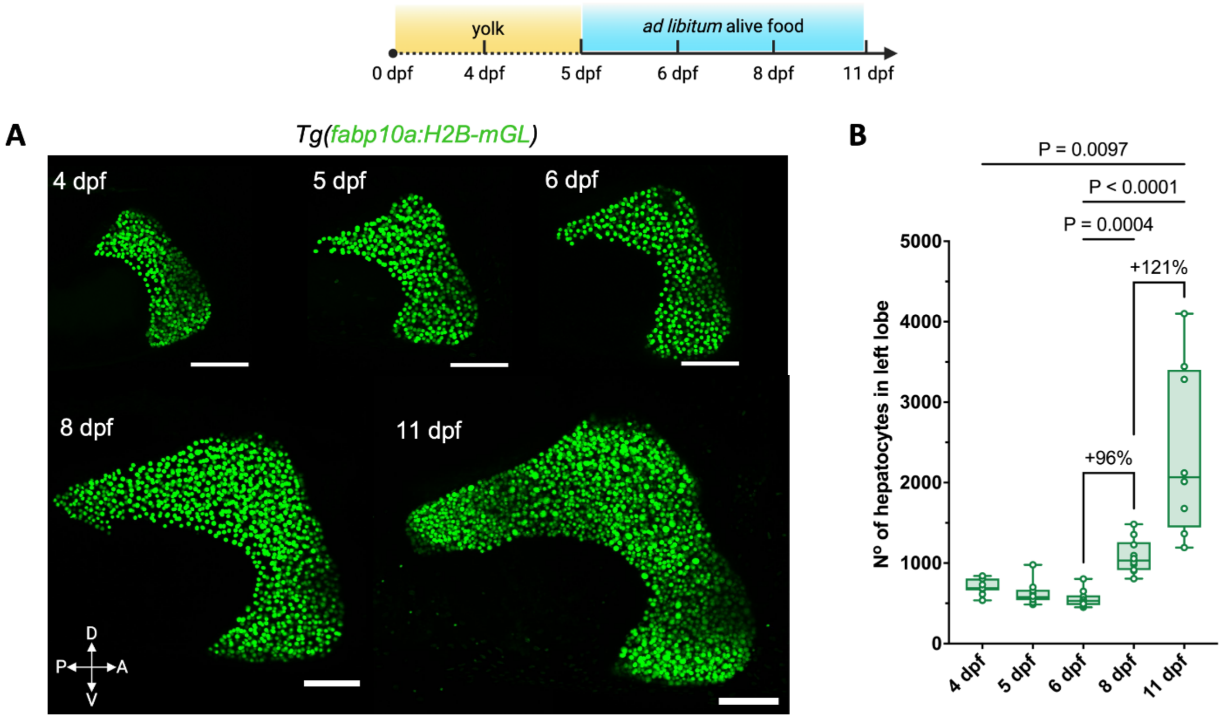

**Figure S1: Rapid increase of hepatocyte population in late larval stage zebrafish. (A)** Until 5 dpf, zebrafish larvae obtain their nutrients from the yolk. From 5 dpf onwards, they are provided with ad libitum live food, specifically rotifers. Live confocal images of the left lobe of *Tg(fabp10a:H2B-mGreenLantern)* zebrafish liver at 4, 5, 6, 8 and 11 dpf. Scale bars: 100 μm. **(B)** Growth curve was assessed by quantifying the number of hepatocytes in the left lobe. Quantification revealed a significant increase in hepatocyte numbers, with a 96% increase observed between 6 and 8 dpf, and a 121% increase between 8 and 11 dpf (Kruskal-Wallis test). These findings underscore the rapid proliferation of hepatocytes during this critical growth period.

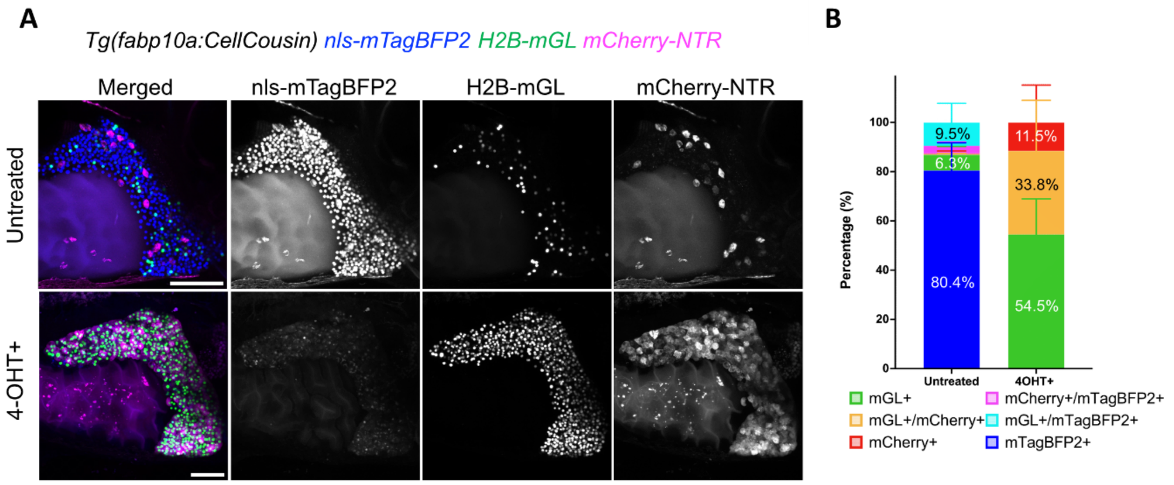

**Figure S2: Highly efficient Cre/Lox recombination in *Tg(fabp10a:CellCousin)* livers.**

**(A)** Live confocal imaging of liver in 9 dpf zebrafish larvae untreated and 4-OHT-treated. In untreated animals, presence of H2B-mGL+ and mCherry+ hepatocytes show the background recombination due to leaky $CreER^{T2}$ recombination driven by the *fabp10a* promoter. 4-OHT-treatment lead to the switching of nls-mTagBFP2 fluorescence label to H2B-mGL (nuclear green) or mCherry (cytoplasmic magenta). **(B)** Mean ± SD of the percentage of hepatocyte labeling in untreated (n=9) and after 4-OHT treatment (n=5). Scale bars: 100 μm (A).

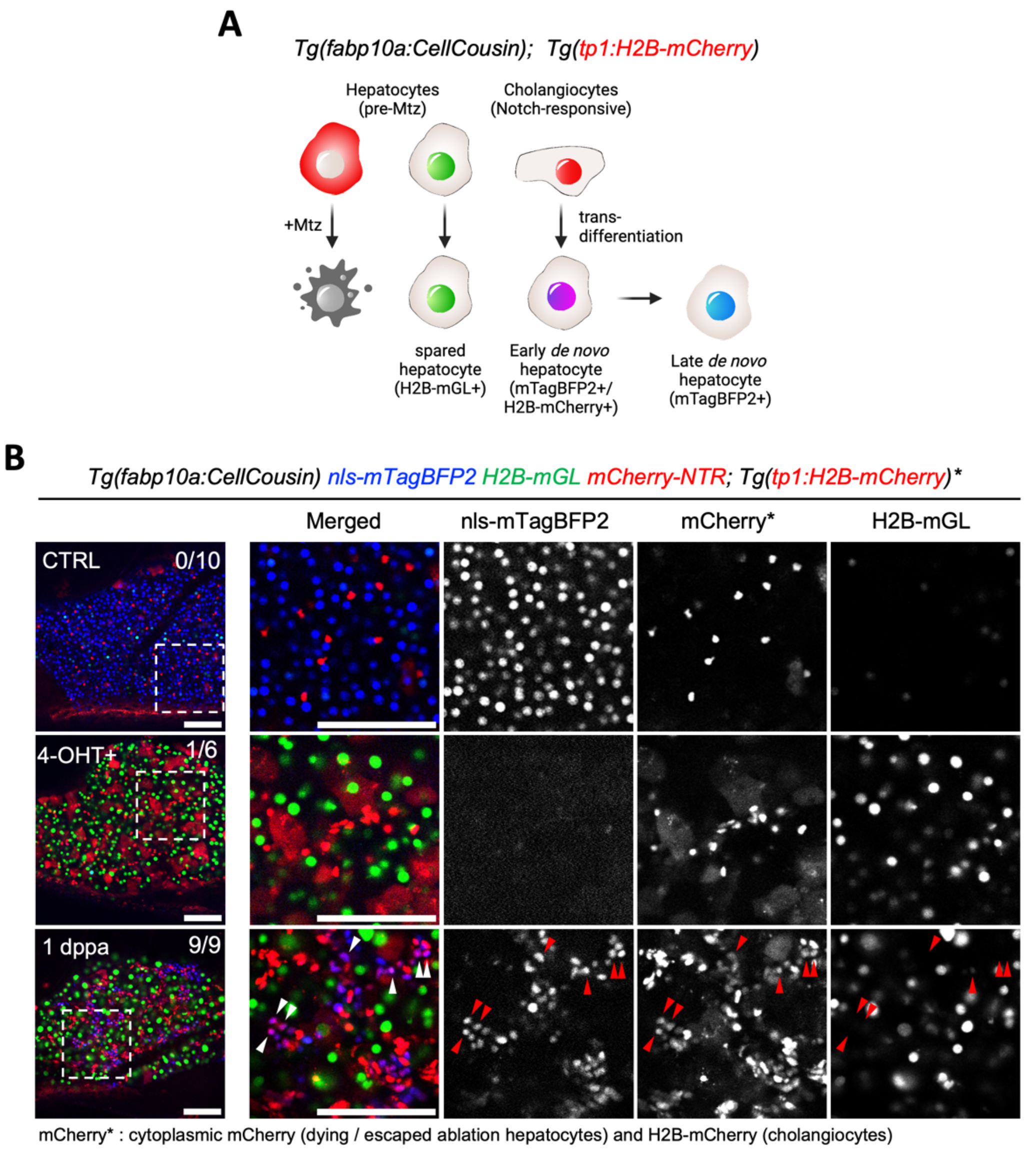

**Figure S3: Cholangiocytes are the source of *de novo* hepatocytes after partial ablation.**

**(A)** Schematic illustrating the lineage tracing of cholangiocytes after partial ablation via the CellCousin method. Hepatocytes are labelled with H2B-mGL+ or mCherry-NTR+ after 4-OHT treatment in the *Tg(fabp10a:CellCousin)* line. Cholangiocytes, which are Notch-responsive, are marked with bright H2B-mCherry+ (nuclear red) using the *Tg(tp1:H2B-mCherry)* line. After MTZ administration, Specific ablation of mCherry-NTR+ hepatocytes leads to a liver with spared H2B-mGL+ hepatocytes and *de novo* nls-mTagBFP2+ hepatocytes. Early *de novo* hepatocytes derived from cholangiocytes are detected as nls-mTagBFP2+/H2B-mCherry+ (nuclear magenta), while late *de novo* hepatocytes, due to dilution of H2B-mCherry label, are marked only with nls-mTagBFP2. **(B)** Confocal images of livers from *Tg(fabp10a:CellCousin); Tg(tp1:H2B-mCherry)* in 13 dpf untreated, 4-OHT-treated and 1 dppa zebrafish larvae using *in vivo* live imaging. nls-mTagBFP2+/H2B-mCherry+ cells were observed in 0 / 10 livers in untreated animals, in 1 / 6 livers in 4-OHT treated animals, and in 9 / 9 livers at 1 dppa after MTZ administration. Scale bars: 100 $\mu$m (B).

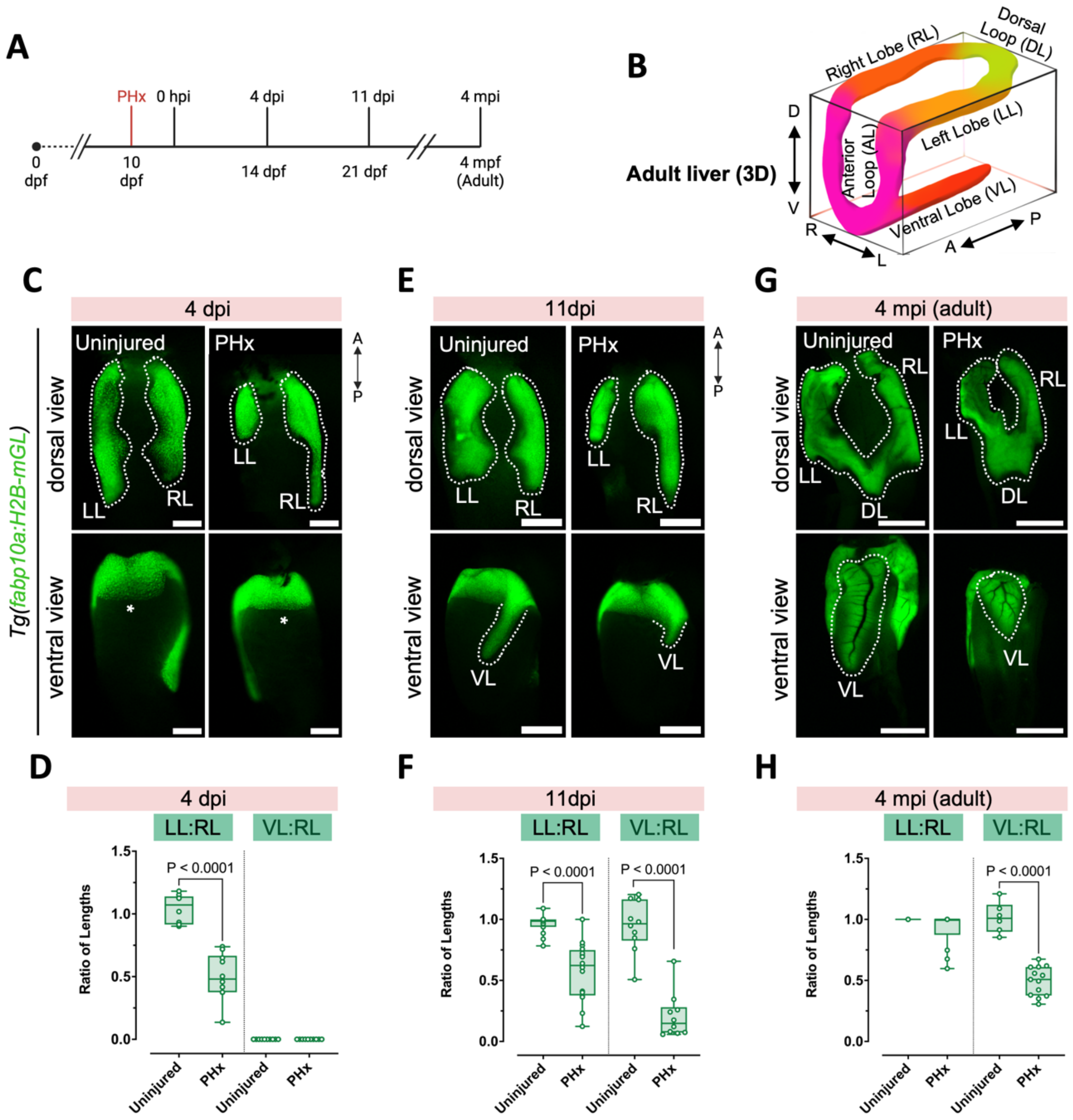

**Figure S4: Morphological recovery of the liver after PHx.**

**(A)** Experimental strategy for assessing regeneration after PHx. **(B)** Schematic illustrating liver lobes of an adult zebrafish. **(C, E, G)** Whole-mount images of liver (green) with dorsal (above) and ventral (below) views (left lobe, LL; right lobe, RL; ventral lobe, VL) at 4 dpi (asterisk: missing VL) (E), at 11 dpi (G), and at 4 mpi (J). **(D, F, H)** Quantifications of the ratio of the lengths of LL to RL and VL to RL at 4 dpi (n=10, each) (D), at 11 dpi (uninjured n=15, PHx n=16) (F), and at 4 mpi (uninjured n=6, PHx n=13) (H) (Mann-Whitney test). Scale bars: 200 μm (C), 500 μm (E), 2 mm (G).

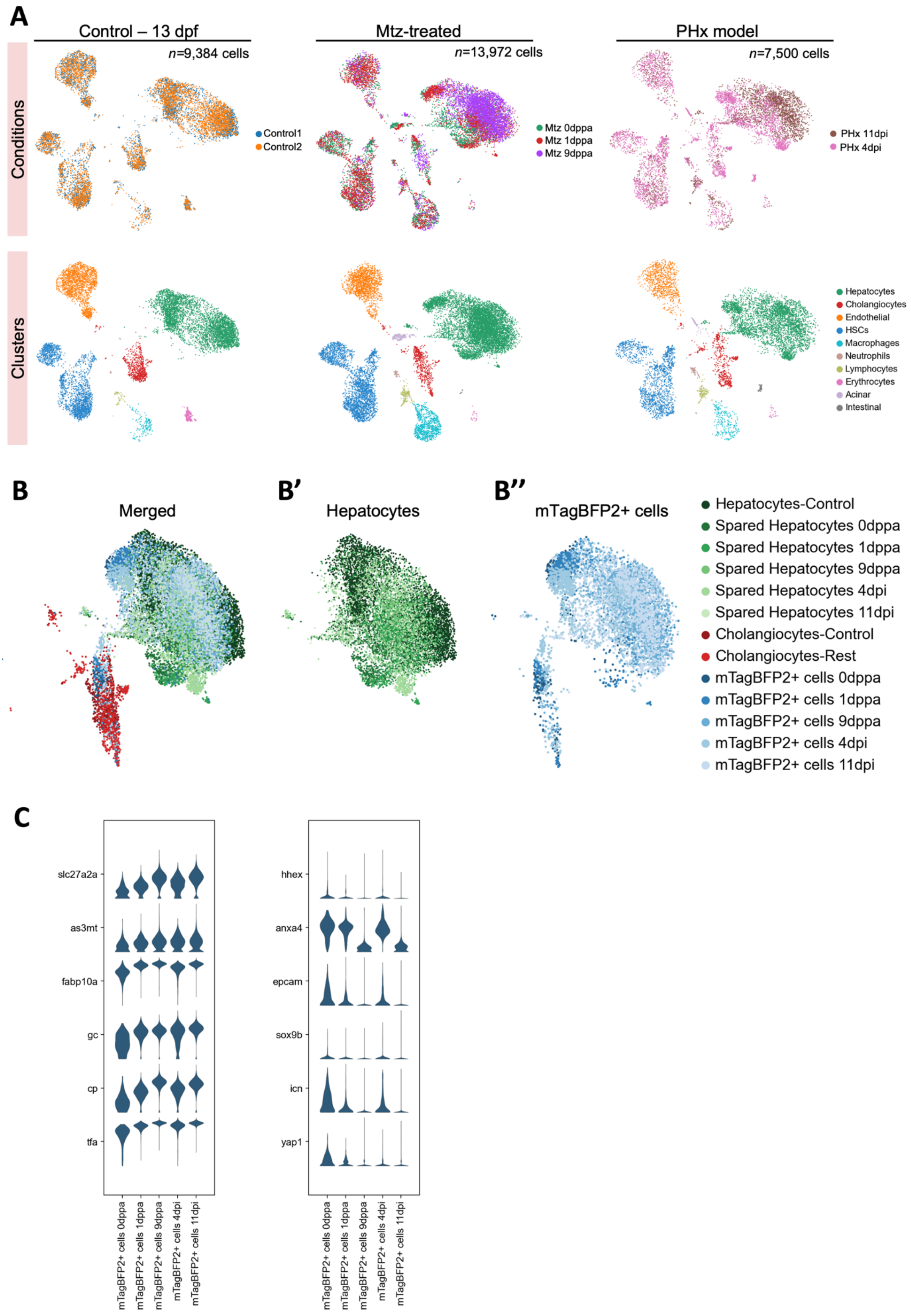

A
Control – 13 dpf
n=9,384 cells
Mtz-treated
n=13,972 cells
PHx model
n=7,500 cells
Conditions
Control1
Control2
Mtz 0dppa
Mtz 1dppa
Mtz 9dppa
PHx 11dpi
PHx 4dpi
Clusters
Hepatocytes
Cholangiocytes
Endothelial
HSCs
Macrophages
Neutrophils
Lymphocytes
Erythrocytes
Acinar
Intestinal
B
Merged
B'
Hepatocytes
B''
mTagBFP2+ cells
Hepatocytes-Control
Spared Hepatocytes 0dppa
Spared Hepatocytes 1dppa
Spared Hepatocytes 9dppa
Spared Hepatocytes 4dpi
Spared Hepatocytes 11dpi
Cholangiocytes-Control
Cholangiocytes-Rest
mTagBFP2+ cells 0dppa
mTagBFP2+ cells 1dppa
mTagBFP2+ cells 9dppa
mTagBFP2+ cells 4dpi
mTagBFP2+ cells 11dpi
C
slc27a2a
as3mt
fabp10a
gc
cp
tfa
hhex
anxa4
epcam
sox9b
icn
yap1
mTagBFP2+ cells 0dppa
mTagBFP2+ cells 1dppa
mTagBFP2+ cells 9dppa
mTagBFP2+ cells 4dpi
mTagBFP2+ cells 11dpi

**Figure S5: Single-cell RNA-sequencing reveals transcriptome of *de novo* hepatocytes following acute injury.**

**(A)** UMAP visualization of zebrafish larval liver cells per condition: uninjured control (left, *n*=9,384 cells), after Mtz treatment (middle, *n*=13,972 cells) and after partial hepatectomy (PHx, *n*=7,500 cells). The top panels show the cells per condition, while the bottom panels display the clusters. **(B)** UMAP highlighting clusters of interest: control cholangiocytes (dark red), cholangiocytes-rest (from injured livers, red), control hepatocytes (dark green) and spared hepatocytes (greens) and mTagBFP2+ cells (blues). Subsets of hepatocyte and mTagBFP2+ cell clusters at different stages after injury are represented with shades of green and blue, respectively. **(C)** Violin plots showing normalized gene expression of hepatocyte-specific markers (left) and cholangiocyte markers (right) in mTagBFP2+ cells across different timepoints in injured conditions.

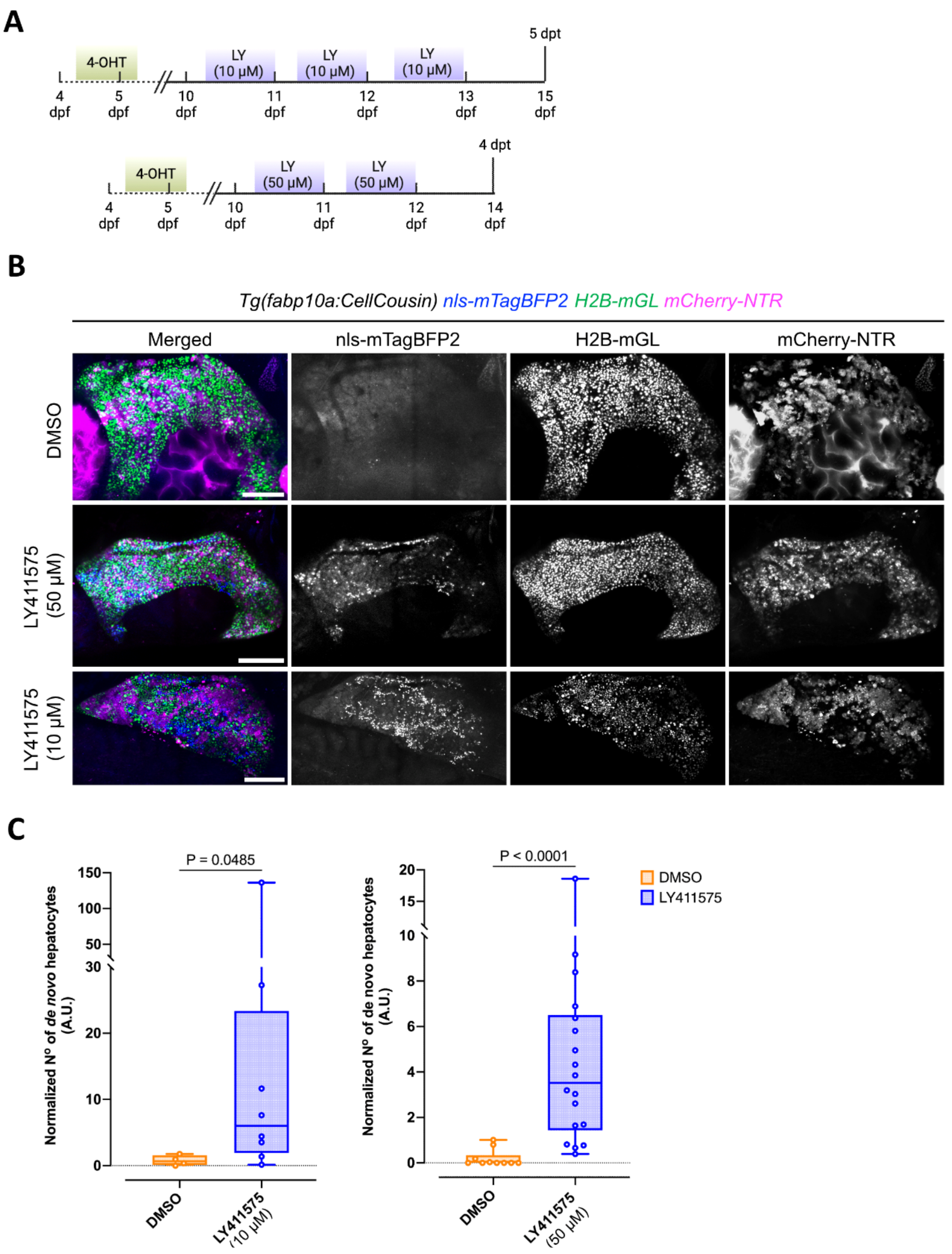

**Figure S6: Inhibition of Notch signalling pathway induces spontaneous transdifferentiation even in the absence of ablation.**

**(A)** Schematic of the experimental strategy for inhibition of Notch signalling pathway in *Tg(fabp10a:CellCousin)* zebrafish larvae. Larvae were treated with 10 μM 4-OHT at 4.5 dpf for 24 hours. To inhibit the Notch pathway, larvae were treated with LY411575

(10 μM or 50 μM) for 16 hours at 3- and 2-days intervals, respectively, and then analysed at 2 days post-treatment (dpt). **(B)** Confocal images of the liver in 15 dpf larvae treated with DMSO, 10 μM, or 50 μM LY411575 reveal the emergence of mTagBFP2+ *de novo* hepatocytes (blue) after LY411575 treatment suggesting spontaneous transdifferentiation from non-hepatocyte sources. Scale bars: 200μm. **(C)** Quantification of normalized mTagBFP2+ *de novo* hepatocytes shows a significant increase following treatment with 10 μM (left, n=8) or 50 μM (right, n=18) LY411575 compared to DMSO controls (n=4 and n=10, respectively) (Mann-Whitney test).

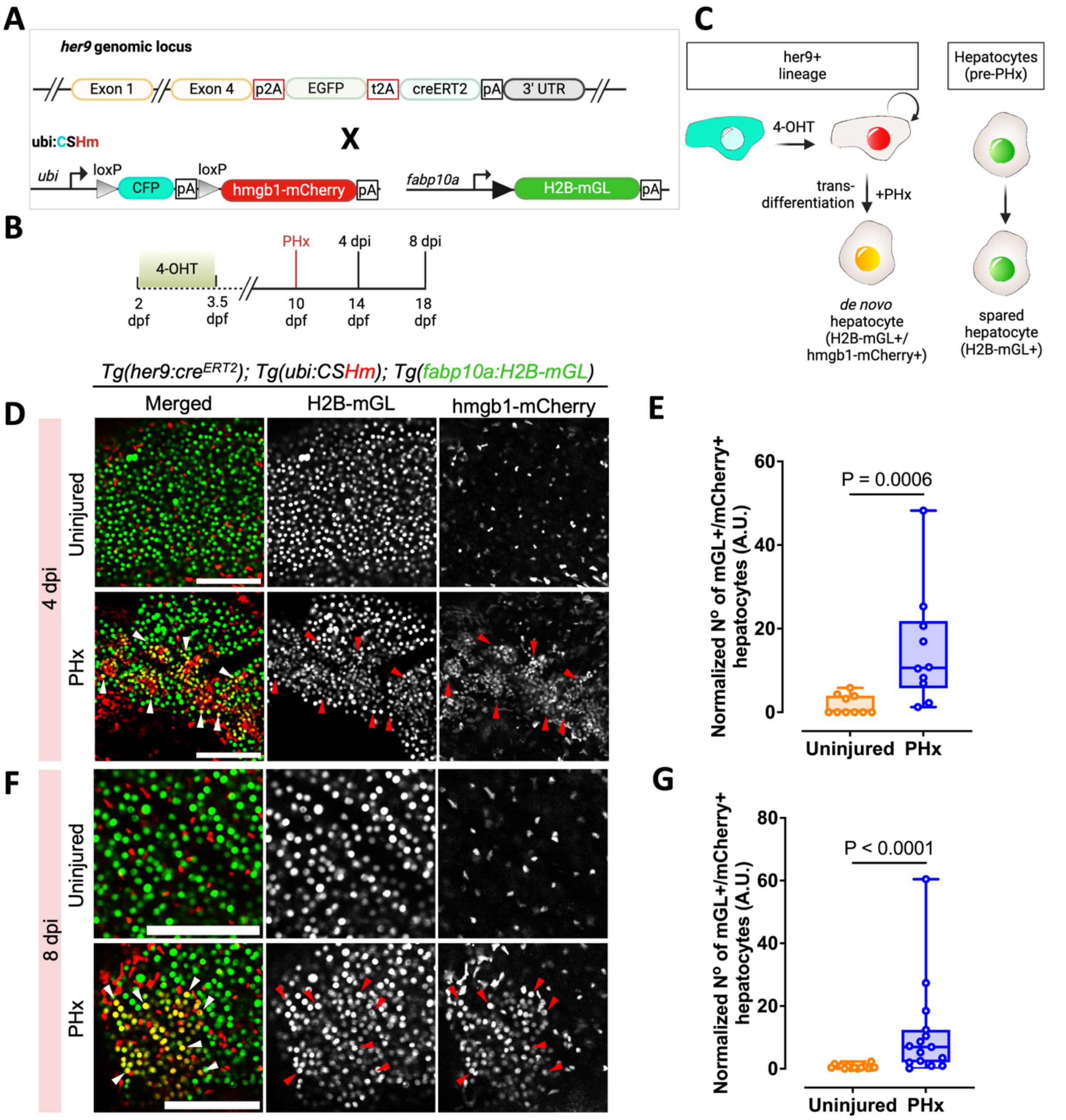

**Figure S7: Lineage tracing strategy to demonstrate the contribution of her9+ cells lineage to *de novo* hepatocytes.**

**(A)** The Cre/loxP strategy used to trace her9-derived *de novo* hepatocytes. Fish carrying a knock-in of p2A-*EGFP-t2A-CreER*$^{T2}$ in the her9 locus are crossed to a ubiquitous Cre-reporter line and a hepatocyte reporter line. **(B)** 4-OHT was administered from 48 hpf to 78 hpf, and the her9 lineage was traced at 4 and 8 dpi following PHx performed at 10 dpf. **(C)** Schematic illustrating the tracing of her9 lineage before and after PHx. her9-expressing cells are labeled with mCherry following 4-OHT treatment. After transdifferentiation into hepatocytes, these cells become H2B-mGL+/mCherry+ as hepatocytes are marked with H2B-mGL. **(D, F)** Confocal images of left lobe in uninjured and PHx conditions at 4 dpi (D) and at 8 dpi (F). Representative H2B-mGL+/mCherry+ hepatocytes are marked with arrowheads. Scale bars: 100 μm. **(E, G)** Quantification of the normalized number of H2B-mGL+/mCherry+ cells representing her9-derived *de novo* hepatocytes at 4 dpi (E) and at 8 dpi (G) shows a significant increase in animals subjected to PHx (n=10, 10, respectively) compared to uninjured animals (n=10, 15, respectively) (Mann-Whitney test).

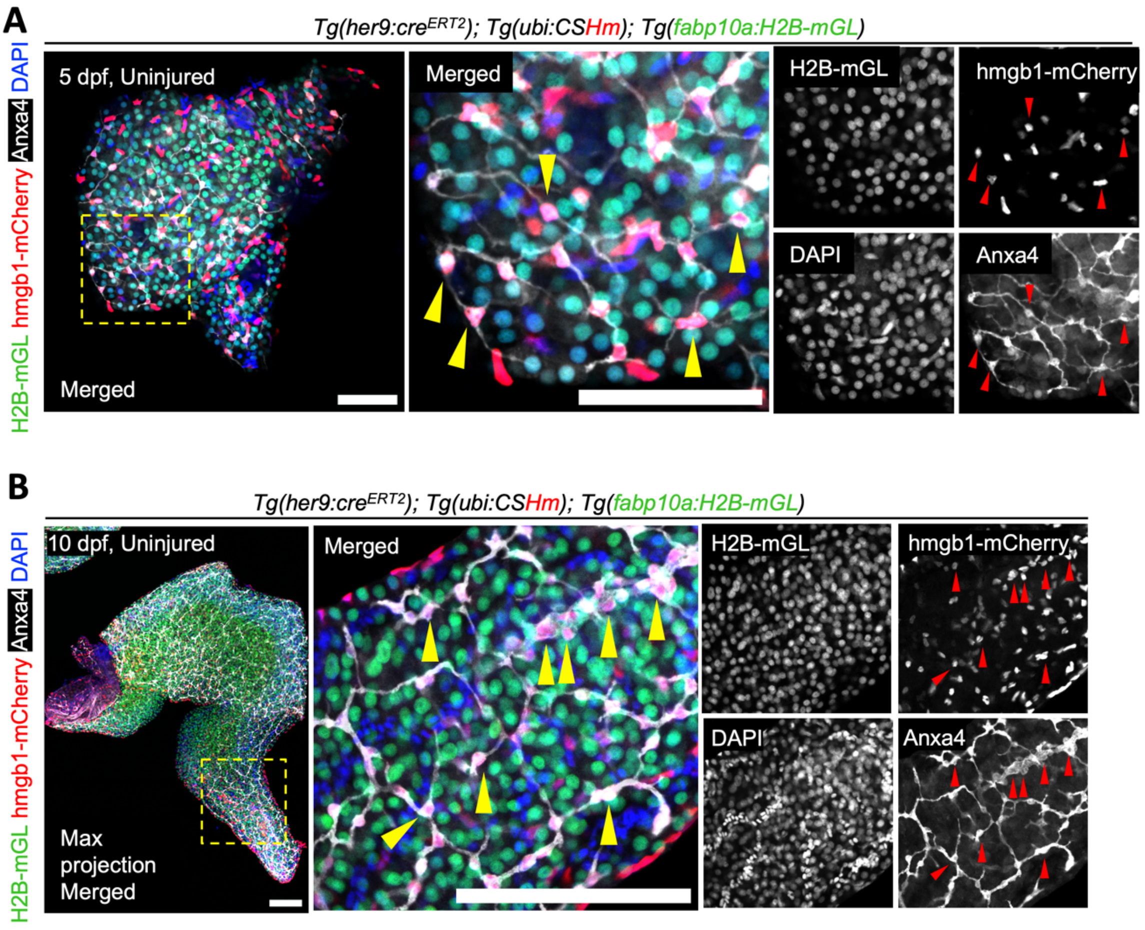

**Figure S8: her9+ cell lineage mainly contains cholangiocytes.**

**(A, B)** The her9 lineage is marked with nuclear mCherry following 4-OHT treatment from 48 to 78 hpf, while hepatocytes are labelled with H2B-mGL. At 5 dpf (A) and at 10 dpf (B), a subset of mCherry cells, but none of the H2B-mGL cells, exhibit Anxa4-immunofluorescence (gray). Anxa4 labelling is specific to cholangiocytes. Representative mCherry cells with Anxa4 label are marked with arrowheads. Scale bars: 50 $\mu$m (A), 100 $\mu$m (B).
